# Deciphering bacteriophage T5 host recognition mechanism and infection trigger

**DOI:** 10.1101/2022.09.22.509047

**Authors:** Séraphine Degroux, Grégory Effantin, Romain Linares, Guy Schoehn, Cécile Breyton

**Affiliations:** Univ. Grenoble Alpes, CNRS, CEA, IBS, F-38000, Grenoble, France

**Keywords:** Bacteriophage, electron cryo-microscopy, host recognition, conformational changes

## Abstract

Bacteriophages, viruses infecting bacteria, recognise their host with high specificity, either binding to saccharide motifs or proteins of the cell wall of their host. In the majority of bacteriophages, this host recognition is performed by Receptor Binding Proteins (RBPs) located at the extremity of a tail. Interaction between the RBPs and the host is the trigger for bacteriophage infection, but the molecular details of the mechanisms are unknown for the majority of bacteriophages. Here, we present the electron cryo-microscopy structure of bacteriophage T5 RBP_pb5_ in complex with its *E. coli* receptor, the iron ferrichrome transporter FhuA. Monomeric RBP_pb5_ is located at the extremity of T5 long flexible tail, and its irreversible binding to FhuA commits T5 to infection. Analysis of RBP_pb5_ structure within the complex, comparison with its AlphaFold2 predicted structure, and its fit into a previously determined map of T5 tail tip in complex with FhuA allow us to propose a mechanism of transmission of RBP_pb5_ receptor binding to the straight fibre, initiating the cascade of events that commits T5 to DNA ejection.

## Introduction

Bacteriophages, bacterial viruses also called phages, are the most abundant and diversified biological entity on Earth (Dion *et al*, 2020). As bacterial killers, but also as a means of genetic material exchange, they shape and drive microbial population evolution and diversity (Suttle, 2007). The vast majority of phages indexed in databases are composed of a capsid protecting the DNA and bear a tail, which can be can be long and contractile (*Myoviridae*), long and flexible (*Siphoviridae*) or short (*Podoviridae*). The tail extremity serves to recognise the host surface, perforate its cell wall and safely channel the DNA from the capsid to the bacterial cytoplasm (Nobrega *et al*, 2018; Linares *et al*, 2020). The distal tip of the tail bears Receptor Binding Proteins (RBPs) that reversibly or irreversibly bind saccharides or proteins. Depending on their morphology, RBPs can be fibres (more than 1000 residues long, requiring chaperones to fold) or more globular RBPs often called Tail Spike Proteins (TSP)(Broeker & Barbirz, 2017). Fibres and TSP of known structure share an elaborately interwoven often trimeric state, rich in β-helices. TSP are multidomain proteins, containing at least a phage binding domain and a receptor binding domain (Goulet *et al*, 2020). The receptor binding domain can also bear enzymatic activity, digesting the saccharide receptors (Broeker & Barbirz, 2017). Because they recognise the bacterial host surface with high specificity and because of their enzymatic activities, phage RBPs can be used to identify bacterial strains for diagnosis (Filik *et al*, 2022), to engineer phages with new host range (Dams *et al*, 2019) or in biotechnology (Santos *et al*, 2018). The stoichiometry of RBP per virion can vary from one copy (*e*.*g*. RBP_pb5_ in colisiphophage T5) to 54 (*e*.*g*. in lactosiphophage TP901-1) and maybe more; phages can also combine RBPs with different binding specificities (Sørensen *et al*, 2021). Multiple binding is proposed to increase affinity of saccharide binding by avidity. It also has been proposed to position the tail tube correctly for cell wall perforation (Taylor *et al*, 2018) and has been shown to trigger infection (*e*.*g*. T4 (Hu *et al*, 2015; Yap *et al*, 2016) and T7 (Hu *et al*, 2013; González-GarcÍa *et al*, 2015)). All RBP structures determined to date are saccharide binding RBPs.

Phage T5, a *Siphoviridae* infecting *E. coli*, is a model phage belonging to the T-series introduced by Delbrück and co-workers in the 1940s (Demerec & Fano, 1945). It bears an icosahedral capsid that protects the 121 kb DNA and a 250-nm long flexible tail (Arnaud *et al*, 2017). At the distal end of the tail is located the host recognition apparatus that consists of three side tail fibres and a central straight fibre. The central fibre is composed of a trimer of the C-terminus part of the Baseplate Hub Protein (or Tal protein) pb3 followed by a trimer of pb4, at the tip of which is found a monomer of the 640-residues RBP, RBP_pb5_ (Heller & Bryniok, 1984; Zivanovic *et al*, 2014; Linares *et al*, 2020). The side tail fibres reversibly bind Lipopolysaccharide (LPS) O-polymannose (Heller & Braun, 1982; Garcia-Doval *et al*, 2015), allowing the phage to ‘walk’ at the surface of the bacterium until RBP_pb5_ irreversibly binds to FhuA, an *E. coli* iron-ferrichrome transporter (Braun, 2009). *Shigella* and *Salmonella* strains also bear FhuA, sharing high identity with *E. coli* FhuA, and at least *Salmonella paratyphi* can also be hosts to T5 (Graham & Stocker, 1977). T5 side tail fibres are however dispensable, as the T5*hd1* mutant that lacks them is only affected in host adsorption rates (Heller & Bryniok, 1984; Saigo, 1978). *In vitro*, T5 mere interaction with purified FhuA commits T5 to infection and triggers DNA ejection (Boulanger *et al*, 1996), making T5 an excellent model to study phage-host interaction at the cellular and molecular level (Arnaud *et al*, 2017; Linares *et al*, 2022). Both RBP_pb5_ and FhuA can be purified and can *in vitro* bind in a 1:1, irreversible complex (Plançon *et al*, 2002; Flayhan *et al*, 2012). Biophysical analysis of the individual FhuA and RBP_pb5_ proteins compared to the complex showed that some secondary structure changes occur in RBP_pb5_ upon binding to FhuA (Flayhan *et al*, 2012). Also, a low resolution envelop of the proteins obtained by SANS (Small Angle Neutron Scattering) did not show large conformational changes in neither FhuA nor RBP_pb5_ upon interaction (Breyton *et al*, 2013), but showed that RBP_pb5_ is an 8 nm long elongated protein. We recently solved T5 tail tip structure before and after interaction with FhuA reconstituted into a nanodisc, and built atomic models for all the tail tip proteins except RBP_pb5_ and FhuA (Linares *et al*, 2022). These structures allowed us to describe the sequence of events underwent by T5 tail tip upon FhuA-RBP_pb5_ binding, namely straight fibre bending, opening of the tail tube, its anchoring to the membrane and finally formation of a transmembrane channel. Thus, unravelling RBP_pb5_ structure before and after interaction with FhuA is the last piece of the puzzle to fully understand the infection trigger mechanism of phage T5 following FhuA binding. This structure is the first one of a non-saccharide binding RBP.

## Results and discussion

### Overall structure of the FhuA-RBP_pb5_ complex

The FhuA-RBP_pb5_ complex being so stable and FhuA having crystallised in different conditions and detergents (*e*.*g*. (Locher *et al*, 1998; Breyton *et al*, 2019)), we actively tried to crystallise it. We could only obtain non-reproducible anisotropically 8-Å diffracting crystals (Flayhan, 2008). To improve the crystals, we produced nanobodies against the complex. This strategy of increasing the hydrophilic domain of membrane proteins has lead previously to success in crystallisation (Desmyter *et al*, 2015). We could isolate four good binders, and submitted the four FhuA-RBP_pb5_-nanobody complexes to crystallisation. In addition to nanobodies, the lanthanide complex XO4 (nucleating and phasing agent (Engilberge *et al*, 2017)) was used to increase the chances of obtaining well-diffracting crystals. In neither condition could reproducibly diffracting crystals be obtained. Reproducible 2-Å diffracting 2D crystals were obtained and analysed in electron diffraction and image analysis in collaboration with Mohamed Chami and Henning Stahlberg (Basel)(Flayhan, 2008). The crystals were however too thick, rendering structure determination too complicated. Finally, taking advantage of the ‘resolution revolution’ in electron cryo-microscopy (cryo-EM)(Kühlbrandt, 2014), we investigated directly the FhuA-RBP_pb5_ structure as a complex in solution by cryo-EM. Even though the complex is relatively small (150 kDa), we obtained a 2.6-Å resolution 3D structure (Figs. 1A, S1, Table S1).

**Figure 1:**
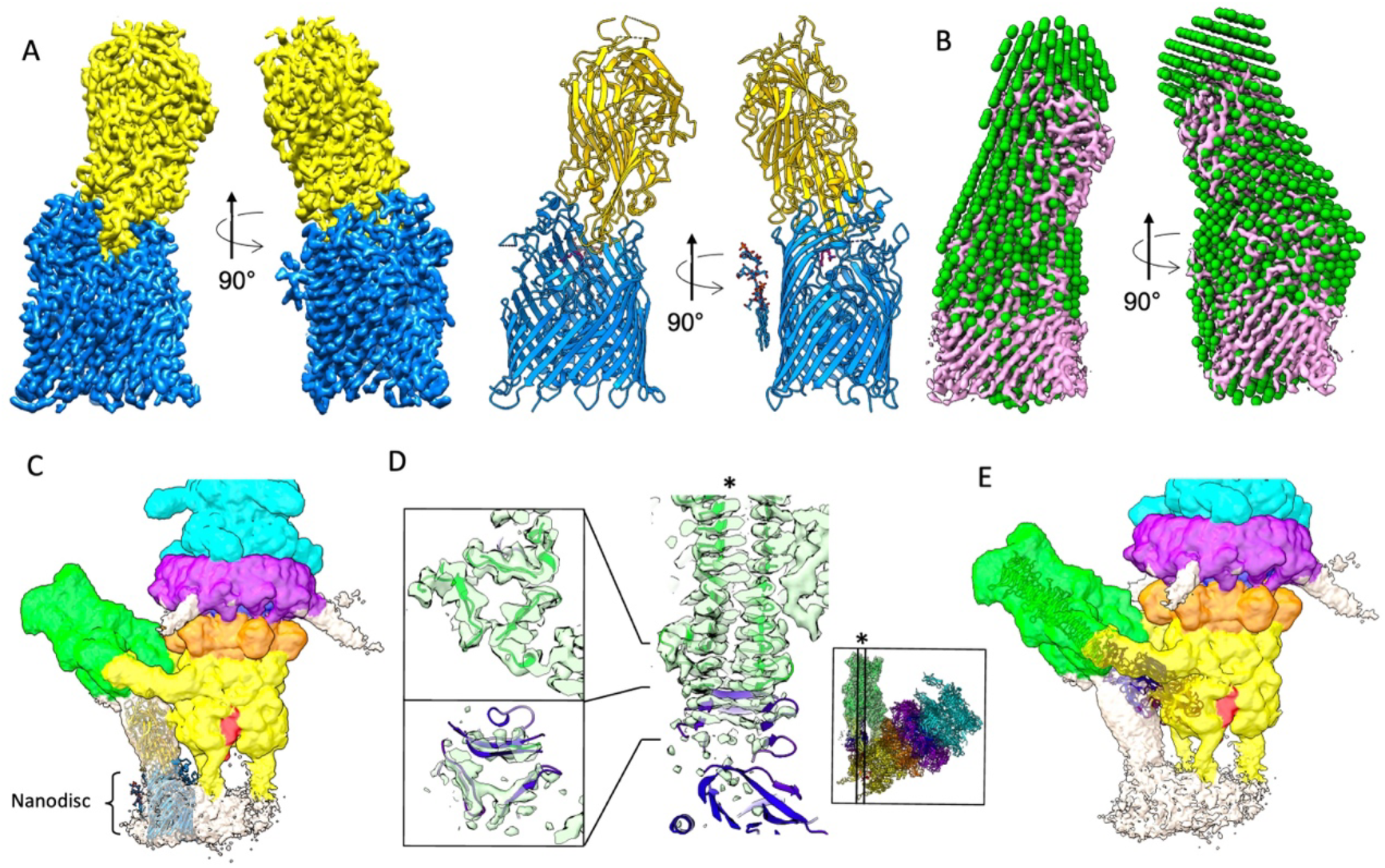
Overall structure of isolated FhuA-RBP_pb5_ and fit in the Tip-FhuA map. **A**. Isosurface view of the FhuA-RBP_pb5_ complex in two 90° related side views (left), and ribbon representation of the modelled proteins with FhuA in blue and RBP_pb5_ in gold (right). **B**. FhuA-RBP_pb5_ map fitted into the SANS envelop of FhuA-RBP_pb5_ (Breyton *et al*, 2013). **C**. Isosurface view at high contour level of the unmasked and unfiltered cryo-EM map of T5 Tip-FhuA (Linares *et al*, 2022) (EMD-14799) with the structure of FhuA-RBP_pb5_ manually fitted into the empty density below pb4 spike and anchoring into the nanodisc. Cyan: density corresponding to pb6, purple: p132, orange: pb9, yellow: pb3, green: pb4 and red: pb2. Unattributed densities are in white. **D**. Manual fit of AF-RBP_pb5_ pb4-binding subdomain into the density extending pb4 spike, with pb4 structure shown in ribbon (EMD-14800, PDB 7ZN4). Top left: Section showing the two last β-strands of pb4 spike trimer, bottom left: section of the density extending pb4 spike. The position of the sections is localised on the longitudinal thin section of pb4 spike map and model (right). The position of the latter section is indicated on the Tip-FhuA model by an * (inset, PDB 7ZN2). **E**. Overall position of AF-RBP_pb5_ in the Tip-FhuA map (same as in C), after manual fitting of pb4-binding subdomain as in D.

The complex and RBP_pb5_ within it have an oblong structure, the interaction surface occurring through the distal end of RBP_pb5_ that penetrates into FhuA barrel down to FhuA plug. The overall FhuA-RBP_pb5_ structure fits well in the SANS envelop (Breyton *et al*, 2013)(Fig. 1B). It also fits well in the map of T5 tail tip interacting with its receptor reconstituted into a nanodisc (Tip-FhuA, Fig. 1C). In the FhuA-RBP_pb5_ structure, both proteins are very well resolved, except for FhuA His_6_-tag located after residue 405 in loop 5 and residues 1-17 including FhuA TonB box. These are disordered in all FhuA structures except when FhuA is crystallised with TonB C-terminal domain (Braun, 2009; Pawelek *et al*, 2006). The lipopolysaccharide that copurifies with FhuA could also be fitted. In RBP_pb5_, the 37 first N-terminal residues, as well as residues 333-352, 365-371 and 427-455 could not be traced in the cryo-EM map. These four sequences point to RBP_pb5_ apex, suggesting, from the fit in Tip-FhuA map, that they form a phage-binding subdomain in interaction with pb4 spike C-terminus. It seems that these loops are thus unstructured without the pb4 spike partner. Unfolding of this small subdomain could also be favoured by the presence of the His-tag in C-terminus of RBP_pb5_, which also points in that direction (Fig. 2A,B).

**Figure 2:**
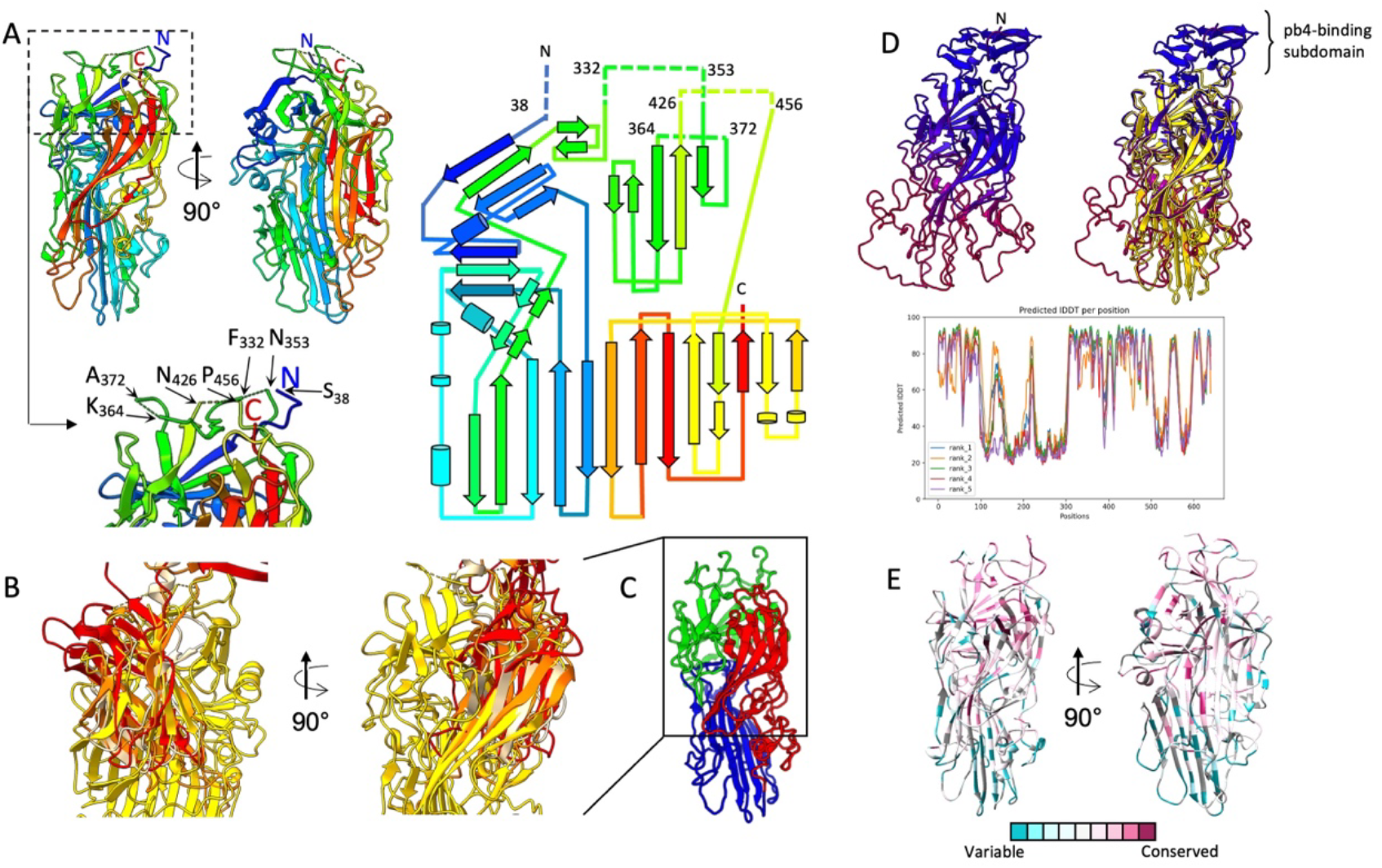
RBP_pb5_ structure analysis. **A**. Top left: ribbon representation of RBP_pb5_ structure in the FhuA-RBP_pb5_ complex context, in two 90° related side views (left) and topology diagram (right), both rainbow coloured (N-terminal: blue, C-terminal: red, indicated)). Bottom left: blow up of the apical region of RBP_pb5_ with indication of the residues bounding the unresolved pb4-binding subdomain. **B**. Dali alignment of RBP_pb5_ (gold) with domains of 5M9F (orange), 1QEX (red), 4A0U (beige). **C**. Subdomain partitioning proposed by the Sword software (https://www.dsimb.inserm.fr/sword/). The position of the aligned subdomain in B is boxed. **D**. Top: best ranked AF-RBP_pb5_ ribbon structure coloured by prediction confidence score (high pLDDT: blue, low pLDDT: dark magenta) alone (left) or aligned with RBP_pb5_ structure within the FhuA-RBP_pb5_ complex (gold)(right). Pb4-binding subdomain is clearly identified at the apex of AF-RBP_pb5_, unresolved in our cryo-EM structure. Bottom: pLDDT scores of the five AF-RBP_pb5_ predictions. **E**. Cryo-EM RBP_pb5_ ribbon representation coloured by sequence similarity / divergence (Ashkenazy *et al*, 2016)(colour key as shown) after ClustalO alignment of a Blast search.

### Analysis of the FhuA-RBP_pb5_ structure

#### RBP_pb5_

As previously proposed (Flayhan *et al*, 2012), RBP_pb5_ folds as a unique domain rich in β-structures. A large and long central 9-stranded β-sheet serves as the spine of the protein. It is surrounded by shorter β-sheets in the apical half and by random coil loops and small α-helices in the distal half of the protein (Fig. 2A). A DALI analysis (Holm, 2020) does not align RBP_pb5_ whole structure with any protein of known structure. However, a β-sandwich subdomain shows structural similarity with other proteins: a close analysis of the aligned DALI domains highlights three proteins with the same topology (Fig. 2B). These three matches are all from phage trimeric proteins, two from RBPs (*S. aureus* phage K gp144, PDB code 5M9F, Dali Z-score = 5.1 and phage T7 gp17, PDB code 4A0U, Dali Z-score = 3.2) and one from a T4 baseplate protein (gp9, PDB code 1QEX, Dali Z-score = 4.9, with however a slight topological difference on the last β-strand). In RBP_pb5_, the β-strands are however longer. This domain displays an original fold, similar however to the knob domain of other viral RBPs, which is believed to interact with the host receptor (Garcia-Doval & van Raaij, 2012). This again illustrates the ‘Lego’ strategy of phages and virus that use and exchange functional modules (Cardarelli *et al*, 2010). RBP_pb5_ structure analysis with the Sword software indeed proposes RBP_pb5_ to be composed of three subdomains, including the common RBP subdomain, as first alternative to a unique domain partitioning (Fig. 2C).

Structure prediction software have done tremendous progress over the last few years (Alexander *et al*, 2021), AlphaFold2 in particular (Jumper *et al*, 2021). To investigate the structure of RBP_pb5_ missing phage-binding subdomain and to compare isolated RBP_pb5_ prediction with our structure, we submitted RBP_pb5_ sequence to AlphaFold2 and obtained five different predicted models (AF-RBP_pb5_)(Fig. 2D). For all models, AF-RBP_pb5_ structure can be divided into two halves: the distal half predicted with a poor confidence (pLDDT < 50)(pLDDT: predicted local-distance difference test), while the apical half is predicted with high confidence (pLDDT > 70). This includes the N-terminus and disordered loops of RBP_pb5_, which are predicted to form a small β-helix, attached to RBP_pb5_ core by five linkers (residues 37-45, 327-336, 353-357, 426-432 and 456-463). This β-helix would extend perfectly pb4 spike: indeed, it fits very well in the density present in Tip-FhuA cryo-EM map extending pb4-β-helix (Fig. 1D). However, the rest of AF-RBP_pb5_ does not fit at all the rest of the low-resolution density available for RBP_pb5_ (Fig. 1E). Indeed, our Tip-FhuA map rather suggests a different angle between pb4 binding subdomain and the rest of RBP_pb5_, made possible by the flexibility given by the five linkers (see also below). This flexibility, visible on negative stain images (see Fig. 5 in Zivanovic *et al*, 2014) explains in part why cryo-EM image analysis of T5 tip and Tip-FhuA did not generate a high resolution map for RBP_pb5_.

Considering the rest of AF-RBP_pb5_ structure prediction, the apical half aligns very well with our RBP_pb5_ structure (rmsd ranging from 0.811 to 0.979 Å over 222 to 291 residues depending on the AF model). This is also coherent with the good level of confidence calculated by AlphaFold2. On the other hand, as expected, alignment of the poorly predicted distal half, corresponding to FhuA binding domain, is pretty bad (overall rmsd ranging from 7.6 to 18.2 Å over all 572 residues for the five AF-RBP_pb5_ models). pLDDT values below 50 are indicated to be strong predictors of disorder (Tunyasuvunakool *et al*, 2021), suggesting that the regions are unstructured in physiological conditions: they would only fold upon interaction with FhuA. This is consistent with biophysical and biochemical data that allowed to conclude that isolated RBP_pb5_ is less structured and sees an increase in its β-sheet content at the expense of other structures, as well as a rigidification of its tertiary structure, upon binding to FhuA (Flayhan *et al*, 2012). This also explains RBP_pb5_ strong thermal stabilisation upon binding to FhuA (Tm = 43°C for RBP_pb5_ alone and 89°C for FhuA-RBP_pb5_)(Flayhan *et al*, 2012).

We also submitted the FhuA-RBP_pb5_ complex to AlphaFold2: the prediction resulted to be far from correct, exhibiting a wrong orientation between FhuA and RBP_pb5_ and a very small interacting surface. RBP_pb5_ does however appear a bit more structured, in particular its large central sheet (Fig. 3D). We have proposed the FhuA-RBP_pb5_ complex to CASP15, let us see the progress they will have made!

**Figure 3:**
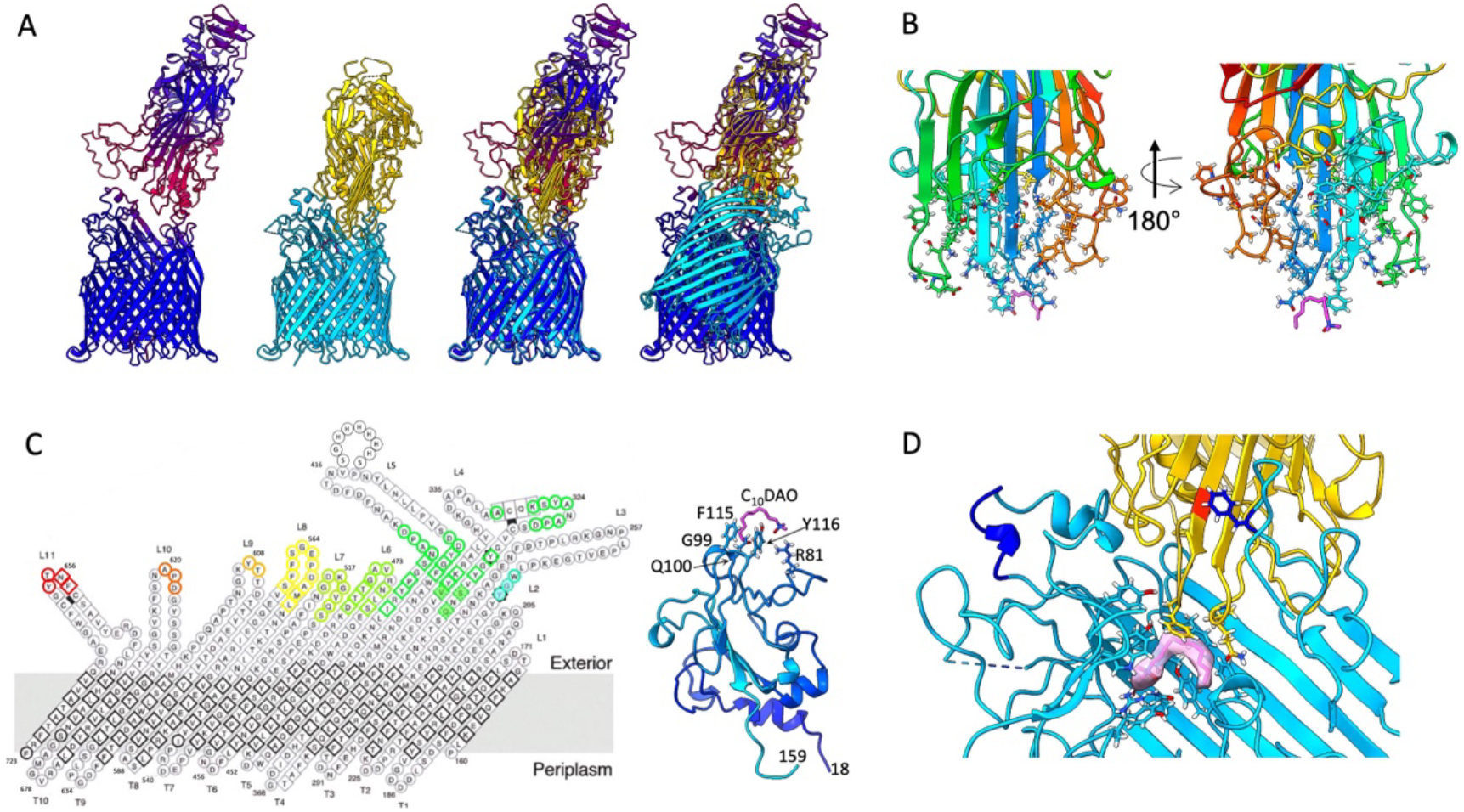
Surface of interaction between FhuA and RBP_pb5_. **A**. From left to right: AF-FhuA-RBP_pb5_ best prediction, coloured by prediction confidence score (high pLDDT: blue, low pLDDT: dark magenta), cryo-EM FhuA-RBP_pb5_ structure (FhuA: cyan, RBP_pb5_: gold), alignment of AF-FhuA-RBP_pb5_ on FhuA and on RBP_pb5_ of the cryo-EM structure. **B**. Two orientations of a rainbow-coloured ribbon representation of RBP_pb5_ distal part, with residues involved in the interactions, as determined by PISA, shown in sticks. The C_10_DAO molecule is shown in pink. **C**. Left: topology diagram of FhuA barrel (from Locher *et al*, 1998), on which are indicated the residues involved in the interaction (dotted line and filled symbols: those interacting exclusively with the C_10_DAO). Right: ribbon diagram of FhuA plug, with residues involved in the interactions and the C_10_DAO molecule shown in sticks. **D**. Zoom on the FhuA-RBP_pb5_ interface. FhuA is in cyan, RBP_pb5_ in gold. Both proteins are in ribbon, with the residues involved in C_10_DAO binding in sticks. The C_10_DAO density is shown in pink. FhuA-PADKGH is highlighted in blue. In RBP_pb5_, G166 is highlighted in orange and FhuA-F566 in blue sticks.

**Figure 4:**
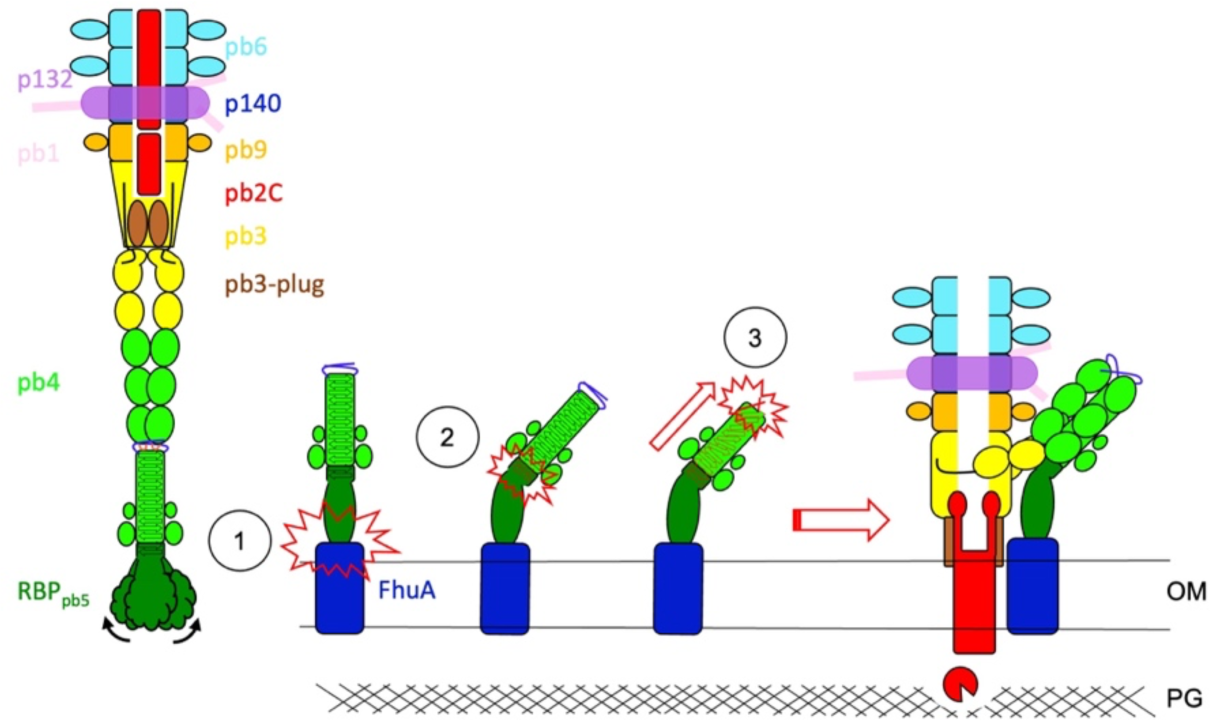
Proposed mechanism of signal transduction of FhuA binding to pb4 spike through RBP_pb5_. Left: within T5 tail, RBP_pb5_ distal half is unstructured and RBP_pb5_ core is flexible with respect to the pb4-binding subdomain. Upon binding to FhuA, RBP_pb5_ distal half gets structured (1). This would induce rigidification of the whole protein, in particular the linkers between the core of RBP_pb5_ and pb4-binding subdomain, which is stabilised at an angle of ∼45° (2). This in turn would induce a modification of the twist of pb4 β-helix (3), which induces the opening of the tail tube, its anchoring to the membrane, the expulsion of the Tape Measure Protein and the formation of a channel across the host outer-membrane (Linares *et al*, 2022).

Whereas there are no RBP_pb5_ homologues in the PDB, a BLAST search fishes a large number of proteins, with identities ranging from 29.0 to 96.3%. They are described as *E. coli, Salmonella, Shigella* or *Klebsiella* phage hypothetical RBPs or RBPs. This is consistent with the fact that the T5-like genus is largely represented in the databases. Clustal alignment and the ConSurf Server (Ashkenazy *et al*, 2016) allow to visualise the distribution of conserved *vs*. variable residues on RBP_pb5_ structure (Fig. 2E). The dichotomy of distribution is quite striking: the apical half of RBP_pb5_ is largely conserved, in particular for the unresolved N-terminus and loops, while the distal half, which is proposed by AlphaFold2 to be largely unstructured when unbound to FhuA and which mediates FhuA interaction, is much more variable. This trend – phage binding domain conserved / receptor binding domain much more variable – is quite general in RBPs, fibres or TSP, highlighting that the receptor binding domains are under heavy evolutionary pressure to diversify as they are in interaction with a large variety of host targets, while the phage binding domain, bearing a structural function, is more constrained by interaction with its phage partner·s (Jonge *et al*, 2019).

#### FhuA

The structure of FhuA has been solved in its apo form, in complex with many ligands and with the C-terminal domain of TonB (Braun, 2009). The different structures are remarkably similar, with rmsd between the eleven different structures below 1.93 Å over all 696 common residues. The main difference between the different structures is the N-terminal: in FhuA apo structures, residues 19-30 are structured as an α-helix, whereas in structures binding TonB-dependant ligands it adopts a conformation which is superimposable to the TonB bound conformation (Braun, 2009). In agreement with the fact that T5 infection is TonB independent, the conformation of RBP_pb5_-bound FhuA residues 19-30 is that of apo FhuA.

### FhuA-RBP_pb5_ interface

RBP_pb5_ interaction with FhuA occurs through the distal β-turns and loops of the central sheet and through random coil loops and small α-helices in the distal half of RBP_pb5_, with a total of 63 residues involved, as calculated by PISA (Krissinel & Henrick, 2004)(Fig. 3A). In FhuA, interactions are mediated by the apical tip of the plug, residues in loops 4-11 and in β-strands 7-12, with a total of 64 residues involved (Fig. 3B). The surface of interaction is large, with a buried area of 2,034 Å_2_, an energy of - 21.4 kcal/mol and it is stabilised by 13 hydrogen bonds and hydrophobic interactions, explaining the quasi-irreversibility of the interaction (Flayhan *et al*, 2012). Interestingly, a C_10_DAO (n-Decyl N,N-dimethylamine N-oxide) molecule, contained in the buffer, could be modelled at the interface between FhuA plug and RBP_pb5_ (Figs. 3C, S1E). It overlaps with ferrichrome hydrophobic binding pocket and interacts, on FhuA side, with 13 residues – four from the plug, four from loops 3 and 4, and five from β-strands 7 to 9, with a hydrogen bond between Y116 and the carboxyl of the aminoxyde head group. The C_10_DAO binding pocket is further completed by four residues from loops of the tip of RBP_pb5_ β-sheet, further stabilising the interaction between the two proteins (Fig. 3C).

T5 is a historic phage and a prototype of siphophages, for which host binding has been extensively studied in the past. The presence of an RBP binding to FhuA has been proposed while studying the *oad* mutant that had a reduced affinity to FhuA (Heller & Bryniok, 1984; Krauel & Heller, 1991). Later, Mondigler *et al*, 1996 identified the *oad* mutation: Gly166 is switched to a Trp. In our structure, Gly166 participates in the binding interface, making C-H…П interaction with FhuA Phe566 (Fig. 3C). A Gly->Trp mutation would thus indeed destabilise the protein-protein interaction interface. Mondigler *et al* also investigated the ability of different RBP_pb5_ deletion and insertion mutants to bind to FhuA. Apart from one (deletion of the 153 C-terminus residues), none retained binding abilities, which is explained by the tight fold of FhuA-bound RBP_pb5_.

FhuA binding to RBP_pb5_ has also been studied from the FhuA perspective: before FhuA structure was known, Killmann *et al*, 1995 investigated T5 binding loops in FhuA by competitive peptide mapping. They identified a peptide – _334_PADKGH_339_ – of FhuA loop 4 that induced DNA ejection when incubated with T5, in a temperature dependant manner. Given the large interaction area and the many loops involved to stabilise RBP_pb5_ bound conformation, it is surprising that interaction of RBP_pb5_ with such a small peptide would induce the large conformational changes resulting in DNA ejection. Furthermore, as seen in our structure, this sequence does not appear to be involved in RBP_pb5_ binding (Fig. 3C). We could not reproduce the results with a slightly longer peptide (APADKGHY). Point mutation impairing T5 binding could not be isolated unless they perturb FhuA overall structure (Braun, 2009) and deletion of loops three to eleven, except for loop 8, did not prevent T5 binding (Endriss & Braun, 2004). This can be explained by the large interacting surface between the two proteins, which would compensate point mutations or loop deletions. There does not seem to be a rational for loop 8 being more important for RBP_pb5_ binding than the other ones. Deletion of this loop also impairs T1 and Φ80 binding, and ferrichrome transport, suggesting its deletion might rather prevent FhuA proper folding. A FhuA plug-less mutant was also investigated for its ability to bind T5: an *E. coli* strain deleted for endogenous FhuA and expressing FhuAΔplug is resistant to T5, thus RBP_pb5_ does not bind to FhuAΔplug (Braun *et al*, 2003). Given the surface of interaction provided by FhuA barrel loops, this result suggests that the latter are either flexible or adopt a different conformation in FhuAΔplug than in WT FhuA, preventing RBP_pb5_ binding. Indeed, the plug does not provide a larger surface of interaction than individual FhuA barrel loops. The role of the plug is however important in the overall stabilisation of the interaction between FhuA and RBP_pb5_, as previously suggested (Flayhan *et al*, 2012): whereas FhuA displays two Tm, one for the plug (64°C) and one for the barrel (75°C), the complex displays a unique Tm which is shifted to 89°C.

T5 is described as infecting *E. coli*; however, some other strains also bear FhuA. Thus, we tested T5 sensitivity on different *E. coli*, Shigella and Salmonella strains, which were chosen because of the different level of FhuA identity with *E. coli* lab strain FhuA, ranging from 96 to 100% (Fig. S2). T5 sensitivity tests were performed in the presence of dipyridyl, and iron chelator that boosts FhuA expression. Surprisingly, only *Salmonnella bongori* is sensitive to T5. FhuA sequence alignment shows however some differences in the residues interacting with RBP_pb5_, confirming the resilience of the FhuA-RBP_pb5_ interaction. We checked T5 irreversible adsorption on all strains, but only *Salmonella bongori* showed adsorption, comparable to that of *E. coli*. Interestingly, *E. coli* EPEC and the *Shigella* strains are resistant to T5 despite no difference in the interacting residues (Fig. S2). Resistance to T5 may be explained by FhuA being blocked by a surperinfection exclusion protein coded by a prophage in their genome or by the synthesis of a capsule, shielding FhuA from the phage.

### Conclusion

We present here the first structure of an RBP, RBP_pb5_, in complex with its protein receptor, FhuA. Analysis of this structure compared to that predicted of RBP_pb5_ and fitted in the T5 tail tip map after interaction with FhuA (Tip-FhuA)(Linares *et al*, 2022) allow us to propose a mechanism that would transmit FhuA-binding to pb4 through RBP_pb5_ (Fig. 5): the apical subdomain of AF-RBP_pb5_ shows a small structured β-helix that fits well in our Tip-FhuA map, in the densities extending pb4 β-helix spike. This RBP_pb5_ subdomain would serve as an adaptor between the trimeric pb4 and the monomeric RBP_pb5_. The presence of linkers between this subdomain and RBP_pb5_ core confers flexibility between these two parts of the protein. AF-RBP_pb5_ predicted structure and biophysical analysis of RBP_pb5_ (Flayhan *et al*, 2012) suggest that the distal half of RBP_pb5_ is unstructured when unbound to FhuA. Upon binding to FhuA, RBP_pb5_ distal half is structured, rigidifying the protein. In our Tip-FhuA map, RBP_pb5_ is stabilized with a ∼45° angle between pb4-binding subdomain and the protein core. We suggest that rigidification of RBP_pb5_ distal half upon FhuA binding is transmitted to the pb4-binding subdomain, imposing a kink within RBP_pb5_. This rigidification and kink would constrain RBP_pb5_ pb4-binding subdomain, in turn constraining pb4 β-helix spike, inducing the different twist in pb4 β-helix that initiates the conformational change domino resulting in tail tube opening (Linares *et al*, 2022). Definite confirmation of this hypothesis will require the structure of pb4-bound RBP_pb5_ before and after interaction with FhuA.

## Materials and Methods

### Bacterial and phage strains

The heat-stable deletion mutant T5st0 was used for sensitivity and adsorption test. *E. coli* F, a fast-adsorbing strain for T5, was used for the production of T5st0 and for control of free T5 in adsorption tests. *Shigella flexneri* (CIP 106171), *Shigella sonnei* (CIP 106204), *Salmonella enterica serovar* Typhimurium (CIP 104474), *Salmonella enterica indica* (CIP 102501T), *Salmonnella bongori* (CIP 82.33T) and *E. coli EPEC* (EntetoPathogen *E. coli*)(CIP 52.170) were used for sensitivity to T5 and adsorption tests. *E. coli* strain AW740 and BL21(DE3) were used for FhuA and RBPpb5 expression respectively.

#### FhuA and RBPpb5 production and purification

FhuA was purified from the *E. coli* strain AW740 transformed with a plasmid encoding the *fhuA* gene in which a His-tag coding sequence has been inserted in extracellular loop L5. RBPpb5 was purified from the *E. coli* strain BL21(DE3) carrying the *oad* gene encoding RBPpb5 fused to a His-tag coding sequence in 3’ of *oad*, in a pET-28 vector (Plançon *et al*, 2002). Cells were grown in LB medium at 37 °C in the presence of 100 µM of the iron-chelating agent dipyridine for AW740 cells and at 20 °C without induction for BL21(DE3) cells. FhuA and RBPpb5 purifications were carried out as described (Flayhan *et al*, 2012). The FhuA-RBPpb5 complex was formed by adding equimolar amounts of the two proteins, which results in 100% complex formation as described in (Plançon *et al*, 2002). Detergent exchange on the FhuA-RBPpb5 complex was performed by a 10 times dilution in deionised water and centrifugation at 100,000 *g* for 45 min at 4°C. The pellet was rinsed with water and resuspended in 20 mM Tris, 1.6% C10DAO at a protein concentration of 4.3 mg/mL.

#### Cryo-EM sample preparation

Typically, 3.5 µL of the FhuA-RBPpb5 complex were deposited on a freshly glow discharged (25 mA, 30 sec) Cu/Rh 400 mesh Quantifoil R 2/1 EM grids and flash-frozen in nitrogen-cooled liquid ethane using a ThermoFisher Mark IV Vitrobot device (100% humidity, 20°C, 2 s blotting time, blot force 1). Preliminary screening of freezing conditions has been performed on an F20 electron microscope.

Two different datasets were acquired on the same grid. For both dataset, 60-frame movies with a total dose of 60 e^-^/Å^2^ were acquired on a ThermoFisher Scientific Titan Krios G3 TEM (European Synchrotron Radiation Facility, Grenoble, France) (Kandiah *et al*, 2019) operated at 300 kV and equipped with a Gatan Quantum LS/967 energy filter (slit width of 20 eV used) coupled to a Gatan K2 summit direct electron detector. Automated data collection was performed using ThermoFisher Scientific EPU software. A nominal magnification of 130.000x was used, resulting in a calibrated pixel size at the specimen level of 1.052 Å/pixel. For the first dataset, 777 movies were acquired with a phase plate, close to focus (between -0.5 and -1.0 µm) while, for the second dataset, 8752 movies were acquired without phase plate and with a defocus ranging between -1.2 and -2.6 µm

#### EM image processing

For both datasets, the processing was done with Relion (Zivanov *et al*, 2018). Motion-correction using 5×5 patches was done with Motioncor2 (Zheng *et al*, 2017) while CTF estimation was done with Gctf (Zhang, 2016).

For dataset 1, an initial set of particles was selected with the Laplacian of Gaussian (LoG) picker in Relion. After 2D classification with 2 times binned particles, nicely defined 2D classes were used as references for a template-based picking. Following another 2D classification, an initial model was obtained in Relion, which was then used to get a first 3D reconstruction. 3D classification (without a mask) was performed to select the best particles which were then used to compute a final 3D reconstruction at 6.7 Å (at FSC = 0.143) into which the atomic model of FhuA (PDB 1QFG) could be un-ambiguously fitted (not shown).

For dataset 2, 2D projections of the 3D model obtained from dataset 1 were used as a template to automatically pick all micrographs. 2,191,586 particles were obtained, binned two times and further subjected to two rounds of 2D classification to remove obvious outliers. A first 3D classification was done without any mask and the selected particles were re-extracted without binning. Following a 3D refinement step, two successive 3D classifications with a mask following the contour of the complex were performed without any image alignment, leading to a homogenous subset of 109,350 particles. Then, particle polishing and two rounds of CTF refinements (magnification anisotropy, beam tilt and per-particle defocus) were performed before a last 3D refinement yielded the final 3D reconstruction at 2.6 Å average resolution (at FSC = 0.143). The final map was sharpened using DeepEMhancer (Sanchez-Garcia *et al*, 2021).

#### Model building

The RBPpb5 protein model was built *de novo* in FhuA-RBPpb5 cryo-EM map using Coot (version 0.9.2)(Morin *et al*, 2013). FhuA was adapted from a FhuA structure solved by X-ray crystallography (PDB 2GRX). The two individual proteins and the complex were then refined using PHENIX (version 1.18.2-3874) Real Space Refine tool (Morin *et al*, 2013). Structure validation was done using the Molprobity online tool.

#### Phage sensitivity tests

*Shigella, Salmonella* and *E. coli* strains were grown in LB medium at 37°C, 180 rpm until an OD600nm of 0.5. 1.2 ml of LB soft agar are inoculated with 300 µL of culture, and spread on an LB Petri dish. Serial dilutions of phage T5, at 0 to 10^9^ PFU/mL, were directly spotted on the soft agar. The titre was estimated by counting lysis plaques in the spotted area after overnight incubation at 37°C.

#### Irreversible phage adsorption tests

To 5 mL of each strain grown in LB medium at 0.2 OD600nm is added 0.01 M CaCl2, 0.04 M MgCl2 and T5 at a multiplicity of infection of 0.5 and incubated at 37°C. 200 µL of infected cells are collected at different times, vigorously vortexed for 10 s and centrifuged 10 min at 15,000 *g*. The supernatant containing free phages (unabsorbed and reversibly bound particles) are titrated on *E*.*coli F* using the double layer agar method described above.

## Acknowledgments

We would like to acknowledge the thesis work of Ali Flayhan and the shorter or longer internship work of Flavien Grégoire, Charles-Adrien Arnaud, Marie-Ange Marrel, Sarra Landri, Emmi Mikkola and Annelise Vermot who tried to optimise RBPpb5 construct or crystallise the complex with nanobodies, and Edine Hammouche who performed some T5 adsorption tests. We also acknowledge Mohamed Chami and Henning Stahlberg (Biozentrum, Basel) for the 2D crystal adventure, and Alain Roussel and Aline Desmyter (Marseille, https://nabgen.org/) for production of nanobodies directed against the FhuA-RBPpb5 complex. We thank Eric Faudry for a kind gift of the *Salmonella typhidium, Shigella flexneri*, and *E. coli* EPEC strain, Daphna Fenel for quality control by negative staining and Emmanuelle Neumann for help with the F20 electron microscope.

This research was funded by the Agence Nationale de la Recherche, grant numbers ANR-16-CE11-0027 and ANR-21-CE11-0023. It used the EM facility at the Grenoble Instruct-ERIC Center (ISBG; UAR 3518 CNRS-CEA-UGA-EMBL) within the Grenoble Partnership for Structural Biology (PSB). IBS platform access was supported by FRISBI (ANR-10-INBS-05-02) and GRAL, a project of the University Grenoble Alpes graduate school (Ecoles Universitaires de Recherche,) CBH-EUR-GS (ANR-17-EURE-0003). The electron microscope facility is supported by the Auvergne-Rhône-Alpes Region, the Fondation pour la Recherche Médicale (FRM), the Fonds FEDER and the GIS-Infrastructures en Biologie Santé et Agronomie (IBiSA). We acknowledge the provision of in-house experimental time from the CM01 facility at the ESRF. IBS acknowledges integration into the Interdisciplinary Research Institute of Grenoble (IRIG, CEA).

## Author Contributions

CB conceived the project. SD prepared the FhuA-RBP_pb5_ complex. GS and RL optimised cryo-grid preparation. GE recorded and processed the cryo-EM data. SD performed T5 sensitivity and adsorption experiments, and built the atomic model with help from RL. CB analysed and interpreted the structure. CB wrote the paper with contributions from GE, GS and SD. All authors contributed to the editing of the manuscript.

### Data Availability

Cryo-EM density maps of FhuA-RBP_pb5_ and the associated atomic coordinate have been respectively deposited in the Electron Microscopy Data Bank (EMDB) and in the Protein Data Bank (PDB) under the following accession codes: PDB 8B14, EMD 15802.

## Supplementary Figures and Tables

**Figure S1:**
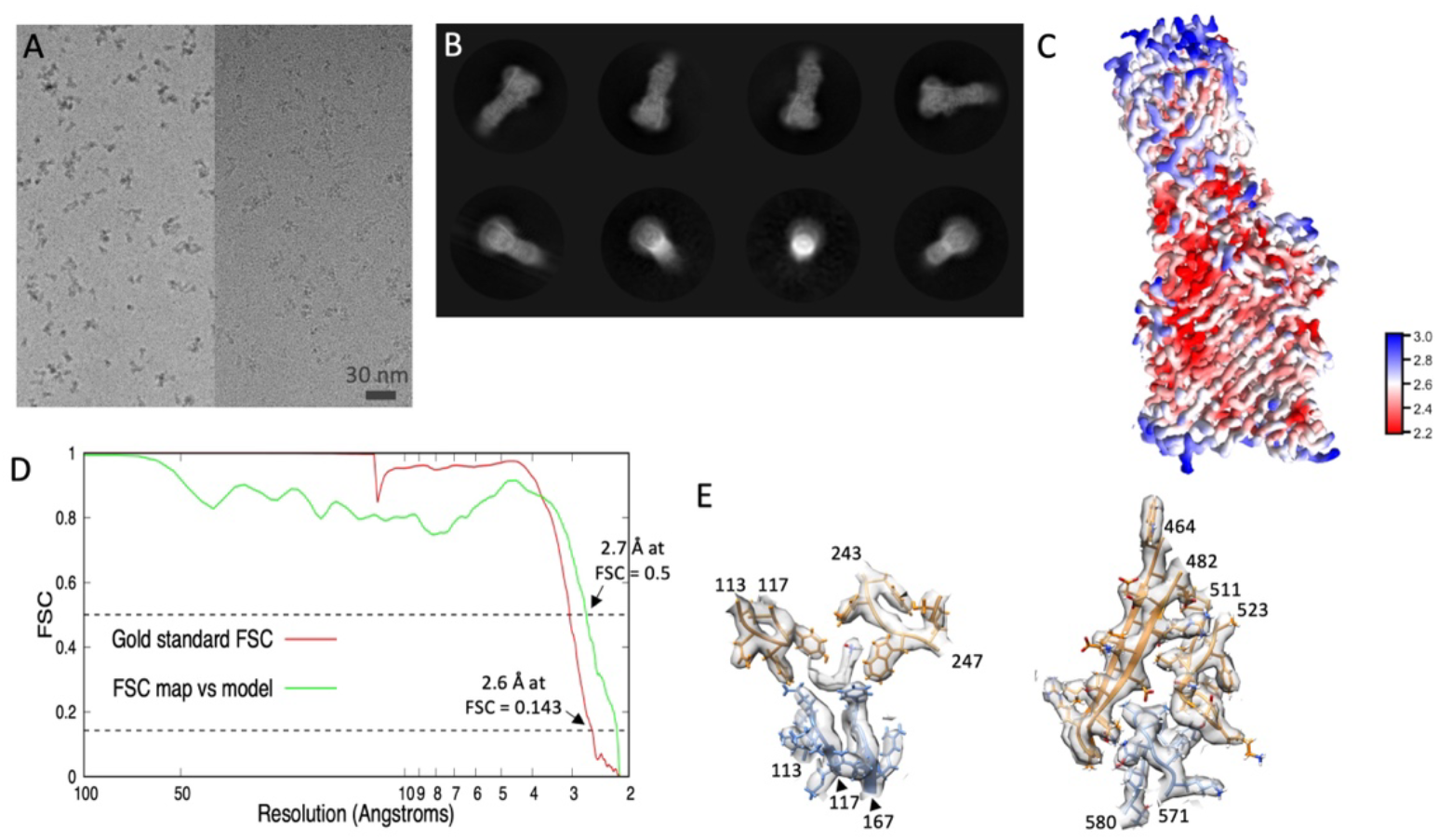
**A**. Cryo-EM field of views of FhuA-RBP_pb5_ molecules with (left) and without (right) phase plate. Scale bar: 30 nm. **B**. Exemplary 2D class averages of FhuA-RBP_pb5_ obtained from the dataset without phase plate. **C**. Local resolution map of the 3D reconstruction of FhuA-RBP_pb5_ obtained with the dataset without phase plate. **D**. Fourier Shell Correlation (FSC) curves: gold standard FSC between two independent 3D reconstructions of FhuA-RBP_pb5_ (red) and FSC between the cryo-EM coulomb potential map and the refined atomic model (green). The two dotted horizontal lines represent FSC = 0.143 and 0.5 which are used as cutoffs to determine the resolutions between the two different sets of maps. **E**. Illustrations of the quality of the obtained 3D reconstruction and atomic model. The coulomb potential map is in transparent gray while FhuA, RBP_pb5_ and C_10_DAO are colored, orange, blue and grey respectively. The left panel is centered around the interaction of FhuA and RBP_pb5_ with C_10_DAO while the right panel represents one of the interacting areas between FhuA and RBP_pb5_ centered around loop 571-580 of RBP_pb5_.

**Figure S2:**
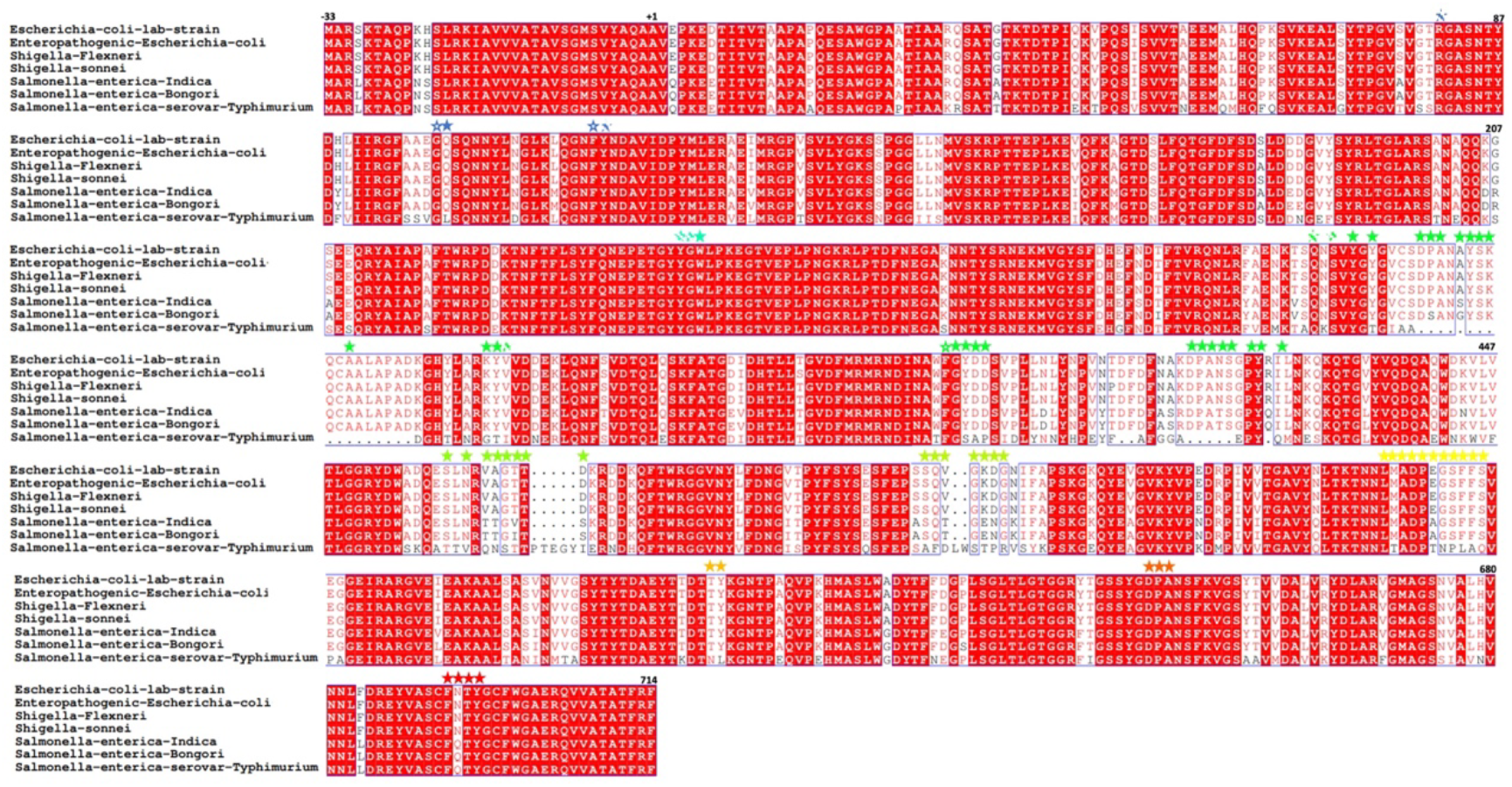
Sequence alignment of FhuA in *E*.*coli, Shigella* and *Salmonella* strains. The alignment was performed with Clustal Omega (https://www.ebi.ac.uk/Tools/msa/clustalo/) and visualised with ESPript (https://espript.ibcp.fr/ESPript/ESPript/). Conserved residues are highlighted in red. Stars show residues involved in the FhuA-RBP_pb5_ interaction. Residues interacting exclusively with RBP_pb5_ (full), with both C10DAO and RBP_pb5_ (empty) or with C_10_DAO only (hatched) are coloured with the same colour code as in Figure 4. Note that the 9 residues of FhuA His-tag are not included in the alignment nor in the numbering.

**Table S1:** Summary of data collection and atomic model statistics

| **Data collection** |  |
| --- | --- |
| Microscope | Titan Krios G3 |
| Voltage (kV) | 300 |
| Magnification | 130,000 x |
| Unbinned pixel size | 1.052 Å/pixel |
| Camera | K2 Summit (Gatan Inc) |
| Exposure time | 8 s |
| Number of frames | 60 |
| Total dose (e<sup>-</sup>/Å<sup>2</sup>) | 60 |
| **Image processing** |  |
| EMDB |  |
| Rotational symmetry | C1 |
| Final number of Particles | 109,350 |
| Average map resolution in Å (FSC 0.143) | 2.6 |
| PDB |  |
| Map to Model resolution in Å (FSC 0.5) | 2.7 |
| Ramachandran favoured (%) | 97.2 |
| Ramachandran outliers (%) | 0.0 |
| Rama Z score | 0.3 |
| Rotamer outliers (%) | 0.8 |
| C-beta deviations | 0 |
| Rms on bond lengths | 0.0072 |
| Rms on bond angles | 1.05 |
| Clashscore | 7.1 |
| Molprobity score | 1.53 |

## References

Alexander LT, Lepore R, Kryshtafovych A, Adamopoulos A, Alahuhta M, Arvin AM, Bomble YJ, Böttcher B, Breyton C, Chiarini V, et al (2021) Target highlights in CASP14: Analysis of models by structure providers. Proteins 89: 1647–1672

Arnaud C-A, Effantin G, Vivès C, Engilberge S, Bacia M, Boulanger P, Girard E, Schoehn G & Breyton C (2017) Bacteriophage T5 tail tube structure suggests a trigger mechanism for Siphoviridae DNA ejection. Nat Commun 8: 1953

Ashkenazy H, Abadi S, Martz E, Chay O, Mayrose I, Pupko T & Ben-Tal N (2016) ConSurf 2016: an improved methodology to estimate and visualize evolutionary conservation in macromolecules. Nucleic Acids Research 44: W344–W350

Boulanger P, le Maire M, Bonhivers M, Dubois S, Desmadril M & Letellier L (1996) Purification and structural and functional characterization of FhuA, a transporter of the Escherichia coli outer membrane. Biochemistry 35: 14216–14224

Braun M, Endriss F, Killmann H & Braun V (2003) In vivo reconstitution of the FhuA transport protein of Escherichia coli K-12. J Bacteriol 185: 5508–5518

Braun V (2009) FhuA (TonA), the career of a protein. J Bacteriol 191: 3431–3436

Breyton C, Flayhan A, Gabel F, Lethier M, Durand G, Boulanger P, Chami M & Ebel C (2013) Assessing the Conformational Changes of pb5, the Receptor-binding Protein of Phage T5, upon Binding to Its Escherichia coli Receptor FhuA. J Biol Chem 288: 30763–30772

Breyton C, Javed W, Vermot A, Arnaud C-A, Hajjar C, Dupuy J, Petit-Hartlein I, Le Roy A, Martel A, Thépaut M, et al (2019) Assemblies of lauryl maltose neopentyl glycol (LMNG) and LMNG-solubilized membrane proteins. Biochim Biophys Acta Biomembr 1861: 939–957

Broeker NK & Barbirz S (2017) Not a barrier but a key: How bacteriophages exploit host’s O-antigen as an essential receptor to initiate infection. Mol Microbiol 105: 353–357

Cardarelli L, Pell LG, Neudecker P, Pirani N, Liu A, Baker LA, Rubinstein JL, Maxwell KL & Davidson AR (2010) Phages have adapted the same protein fold to fulfill multiple functions in virion assembly. Proc Natl Acad Sci U S A 107: 14384–14389

Dams D, Brøndsted L, Drulis-Kawa Z & Briers Y (2019) Engineering of receptor-binding proteins in bacteriophages and phage tail-like bacteriocins. Biochemical Society Transactions 47: 449–460

Demerec M & Fano U (1945) Bacteriophage-Resistant Mutants in Escherichia Coli. Genetics 30: 119–136

Desmyter A, Spinelli S, Roussel A & Cambillau C (2015) Camelid nanobodies: killing two birds with one stone. Curr Opin Struct Biol 32: 1–8

Dion MB, Oechslin F & Moineau S (2020) Phage diversity, genomics and phylogeny. Nat Rev Microbiol 18: 125–138

Endriss F & Braun V (2004) Loop deletions indicate regions important for FhuA transport and receptor functions in Escherichia coli. J Bacteriol 186: 4818–4823

Engilberge S, Riobé F, Pietro SD, Lassalle L, Coquelle N, Arnaud C-A, Pitrat D, Mulatier J-C, Madern D, Breyton C, et al (2017) Crystallophore: a versatile lanthanide complex for protein crystallography combining nucleating effects, phasing properties, and luminescence. Chem Sci 8: 5909–5917

Filik K, Szermer-Olearnik B, Oleksy S, Brykała J & Brzozowska E (2022) Bacteriophage Tail Proteins as a Tool for Bacterial Pathogen Recognition-A Literature Review. Antibiotics (Basel) 11: 555

Flayhan A (2008) Etude strctutale du complexe FhuA-pb5Δ1

Flayhan A, Wien F, Paternostre M, Boulanger P & Breyton C (2012) New insights into pb5, the receptor binding protein of bacteriophage T5, and its interaction with its Escherichia coli receptor FhuA. Biochimie 94: 1982–1989

Garcia-Doval C, Castón JR, Luque D, Granell M, Otero JM, Llamas-Saiz AL, Renouard M, Boulanger P & van Raaij MJ (2015) Structure of the Receptor-Binding Carboxy-Terminal Domain of the Bacteriophage T5 L-Shaped Tail Fibre with and without Its Intra-Molecular Chaperone. Viruses 7: 6424–6440

Garcia-Doval C & van Raaij MJ (2012) Structure of the receptor-binding carboxy-terminal domain of bacteriophage T7 tail fibers. Proc Natl Acad Sci USA 109: 9390–9395

González-GarcÍa VA, Pulido-Cid M, Garcia-Doval C, Bocanegra R, van Raaij MJ, MartÍn-Benito J, Cuervo A & Carrascosa JL (2015) Conformational changes leading to T7 DNA delivery upon interaction with the bacterial receptor. J Biol Chem 290: 10038–10044

Goulet A, Spinelli S, Mahony J & Cambillau C (2020) Conserved and Diverse Traits of Adhesion Devices from Siphoviridae Recognizing Proteinaceous or Saccharidic Receptors. Viruses 12

Graham AC & Stocker BA (1977) Genetics of sensitivity of Salmonella species to colicin M and bacteriophages T5, T1, and ES18. J Bacteriol 130: 1214–1223

Heller K & Braun V (1982) Polymannose O-antigens of Escherichia coli, the binding sites for the reversible adsorption of bacteriophage T5+ via the L-shaped tail fibers. J Virol 41: 222–227

Heller KJ & Bryniok D (1984) O antigen-dependent mutant of bacteriophage T5. J Virol 49: 20–5

Holm L (2020) DALI and the persistence of protein shape. Protein Sci 29: 128–140

Hu B, Margolin W, Molineux IJ & Liu J (2013) The bacteriophage t7 virion undergoes extensive structural remodeling during infection. Science 339: 576–579

Hu B, Margolin W, Molineux IJ & Liu J (2015) Structural remodeling of bacteriophage T4 and host membranes during infection initiation. Proc Natl Acad Sci USA 112: E4919–4928

Jonge PA de, Nobrega FL, Brouns SJJ & Dutilh BE (2019) Molecular and Evolutionary Determinants of Bacteriophage Host Range. Trends in Microbiology 27: 51–63

Jumper J, Evans R, Pritzel A, Green T, Figurnov M, Ronneberger O, Tunyasuvunakool K, Bates R, ŽÍdek A, Potapenko A, et al (2021) Highly accurate protein structure prediction with AlphaFold. Nature 596: 583–589

Kandiah E, Giraud T, de Maria Antolinos A, Dobias F, Effantin G, Flot D, Hons M, Schoehn G, Susini J, Svensson O, et al (2019) CM01: a facility for cryo-electron microscopy at the European Synchrotron. Acta Crystallogr D Struct Biol 75: 528–535

Killmann H, Videnov G, Jung G, Schwarz H & Braun V (1995) Identification of receptor binding sites by competitive peptide mapping: phages T1, T5, and phi 80 and colicin M bind to the gating loop of FhuA. J Bacteriol 177: 694–8

Krauel V & Heller KJ (1991) Cloning, sequencing, and recombinational analysis with bacteriophage BF23 of the bacteriophage T5 oad gene encoding the receptor-binding protein. J Bacteriol 173: 1287–97

Krissinel E & Henrick K (2004) Secondary-structure matching (SSM), a new tool for fast protein structure alignment in three dimensions. Acta Crystallogr D Biol Crystallogr 60: 2256–2268

Kühlbrandt W (2014) Biochemistry. The resolution revolution. Science 343: 1443–1444

Linares R, Arnaud C-A, Degroux S, Schoehn G & Breyton C (2020) Structure, function and assembly of the long, flexible tail of siphophages. Curr Opin Virol 45: 34–42

Linares R, Arnaud C-A, Effantin G, Darnault C, Epalle NH, Erba EB, Schoehn G & Breyton C (2022) Structural basis of bacteriophage T5 infection trigger and E. coli cell wall perforation. 2022.09.20.507954 doi:10.1101/2022.09.20.507954 [PREPRINT]

Locher KP, Rees B, Koebnik R, Mitschler A, Moulinier L, Rosenbusch JP & Moras D (1998) Transmembrane signaling across the ligand-gated FhuA receptor: crystal structures of free and ferrichrome-bound states reveal allosteric changes. Cell 95: 771–8

Mondigler M, Holz T & Heller KJ (1996) Identification of the receptor-binding regions of pb5 proteins of bacteriophages T5 and BF23. Virology 219: 19–28

Morin A, Eisenbraun B, Key J, Sanschagrin PC, Timony MA, Ottaviano M & Sliz P (2013) Collaboration gets the most out of software. Elife 2: e01456

Nobrega FL, Vlot M, de Jonge PA, Dreesens LL, Beaumont HJE, Lavigne R, Dutilh BE & Brouns SJJ (2018) Targeting mechanisms of tailed bacteriophages. Nat Rev Microbiol 16: 760–773

Pawelek PD, Croteau N, Ng-Thow-Hing C, Khursigara CM, Moiseeva N, Allaire M & Coulton JW (2006) Structure of TonB in complex with FhuA, E. coli outer membrane receptor. Science 312: 1399–1402

Plançon L, Janmot C, le Maire M, Desmadril M, Bonhivers M, Letellier L & Boulanger P (2002) Characterization of a high-affinity complex between the bacterial outer membrane protein FhuA and the phage T5 protein pb5. J Mol Biol 318: 557–569

Saigo K (1978) Isolation of high-density mutants and identification of nonessential structural proteins in bacteriophage T5; dispensability of L-shaped tail fibers and a secondary major head protein. Virology 85: 422–433

Sanchez-Garcia R, Gomez-Blanco J, Cuervo A, Carazo JM, Sorzano COS & Vargas J (2021) DeepEMhancer: a deep learning solution for cryo-EM volume post-processing. Commun Biol 4: 874

Santos SB, Costa AR, Carvalho C, Nóbrega FL & Azeredo J (2018) Exploiting Bacteriophage Proteomes: The Hidden Biotechnological Potential. Trends Biotechnol 36: 966–984

Sørensen AN, Woudstra C, Sørensen MCH & Brøndsted L (2021) Subtypes of tail spike proteins predicts the host range of Ackermannviridae phages. Comput Struct Biotechnol J 19: 4854–4867

Suttle CA (2007) Marine viruses--major players in the global ecosystem. Nat Rev Microbiol 5: 801–812

Taylor NMI, van Raaij MJ & Leiman PG (2018) Contractile injection systems of bacteriophages and related systems. Mol Microbiol 108: 6–15

Tunyasuvunakool K, Adler J, Wu Z, Green T, Zielinski M, ŽÍdek A, Bridgland A, Cowie A, Meyer C, Laydon A, et al (2021) Highly accurate protein structure prediction for the human proteome. Nature 596: 590–596

Yap ML, Klose T, Arisaka F, Speir JA, Veesler D, Fokine A & Rossmann MG (2016) Role of bacteriophage T4 baseplate in regulating assembly and infection. Proc Natl Acad Sci USA 113: 2654–2659

Zhang K (2016) Gctf: Real-time CTF determination and correction. Journal of Structural Biology 193: 1–12

Zheng W, Wang F, Taylor NMI, Guerrero-Ferreira RC, Leiman PG & Egelman EH (2017) Refined Cryo-EM Structure of the T4 Tail Tube: Exploring the Lowest Dose Limit. Structure 25: 1436-1441.e2

Zivanov J, Nakane T, Forsberg BO, Kimanius D, Hagen WJ, Lindahl E & Scheres SH (2018) New tools for automated high-resolution cryo-EM structure determination in RELION-3. eLife 7: e42166

Zivanovic Y, Confalonieri F, Ponchon L, Lurz R, Chami M, Flayhan A, Renouard M, Huet A, Decottignies P, Davidson AR, et al (2014) Insights into bacteriophage T5 structure from analysis of its morphogenesis genes and protein components. J Virol 88: 1162–1174

